# Centrin-anchored hydrodynamic shape changes underpin active nuclear rerouting in branched hyphae of an oomycete pathogen

**DOI:** 10.1101/652255

**Authors:** Edouard Evangelisti, Liron Shenhav, Temur Yunusov, Marie Le Naour-Vernet, Philipp Rink, Sebastian Schornack

## Abstract

Polyenergid fungi and oomycetes are phylogenetically distant but structurally similar. To address whether they share similar nuclear dynamics we carried out time-lapse imaging of fluorescently labelled *Phytophthora palmivora* nuclei. Nuclei underwent coordinated bi-directional movements during plant infection. Within hyphal networks growing *in planta* or in axenic culture nuclei are either dragged passively with the cytoplasm or actively become rerouted toward nuclei-depleted hyphal sections and often display a very stretched shape. Benomyl-induced depolymerization of microtubules reduced active movements and the occurrence of stretched nuclei. A centrosome protein localized at the leading end of stretched nuclei, suggesting that like fungi astral microtubule-guided movements contribute to nuclear distribution within oomycete hyphae. The remarkable hydrodynamic shape adaptations of *Phytophthora* nuclei contrast those in fungi and likely enable them to migrate over longer distances. Therefore, our work summarises mechanisms which enable a near equal nuclear distribution in an oomycete. We provide a basis for computational modelling of hydrodynamic nuclear deformation within branched tubular networks.

## Introduction

Although nuclei are usually depicted as immobile, their movement is essential for the growth and development of all eukaryotes, including those with filamentous morphology (Morris, 2000). The nuclear dynamics of fungi have been studied during development in axenic culture as well as during infection of susceptible hosts (Xiang, 2018). Nuclear movements may also contribute to infection success (Vaneault-Fourrey et al., 2006; Fernandez et al., 2014; Jeon et al., 2014; Jones et al., 2016). Recent work in *Magnaporthe oryzae* documented the migration of nuclei into the appressorium and, later into the primary infectious hypha upon rice infection (Fernandez et al., 2014).

Fungal nuclei often distribute equally within hyphae, such as in *Aspergillus nidulans* germ tubes upon asexual spore germination (Morris, 1975). This involves microtubules, cytoplasmic dynein and components of the dynactin complex. Indeed, treatment with the microtubule-depolymerizing drug benomyl or mutations in β-tubulin alter nuclear distribution in *A. nidulans* (Oakley and Morris, 1980). Similar nuclear distribution-defects are observed when *Neurospora crassa* cytoplasmic dynein genes (Xiang et al., 1994) or the dynactin component *Arp1* (Plamann et al., 1994) are mutated. Nuclear distribution is also affected in the *Ashbya gossypii* dynein null mutant, but nuclei accumulate at hyphal tips rather than at the spore end (Alberti-Segui et al., 2001).

Studies in budding yeast (*Saccharomyces cerevisiae*) showed that proper orientation of the mitotic spindle during anaphase is crucial for the mother cell and the bud to receive a single nucleus after nuclear division (Palmer et al., 1992). Such orientation is achieved through astral microtubules (Palmer et al., 1992). Assembly of a functional mitotic spindle also requires precise coordination between the nuclear cycle and the major microtubule organizing centre, also known as the centrosome (Leo et al., 2012). A canonical centrosome is composed of two centrioles surrounded by an electron-dense protein-containing matrix (pericentriolar matrix), which nucleates microtubules (Ito and Bettencourt-Dias, 2018). All fungi except Chytridiomycota possess an acentriolar centrosome termed the spindle pole body (SPB) (Powell, 1980; Ito and Bettencourt-Dias, 2018). The SPB is embedded in the nuclear envelope and nucleates spindle microtubules inside the nucleus as well as astral microtubules toward the cytoplasmic side (Ito and Bettencourt-Dias, 2018). The growing ends of microtubules are occupied by dynein. In *A. gossypii*, dynein capture by the cortical nuclear migration protein Num1 initiates the pulling of cytoplasmic microtubules, thereby moving the attached nucleus (Gibeaux et al., 2017). Orthologs of *Num1* include the *AND1* gene from *M. oryzae* (Jeon et al., 2014) and *ApsA*/*ApsB* from *A. nidulans* (Veith et al., 2005).

Oomycetes are a group of microorganisms with filamentous morphologies that are phylogenetically distant from fungi. They differ from fungi by structural, biochemical and genetic features such as genome size, ploidy, cell wall composition, pigmentation, secondary metabolites and the chemical nature of mating hormones and reserve compounds (Judel-son and Blanco, 2005). Plant-pathogenic oomycetes cause devastating diseases worldwide that impact crop yield, threaten food security and damage natural ecosystems (Fisher et al., 2012; Bebber and Gurr, 2015). For instance, blight, canker and rot diseases caused by plant-infecting oomycetes from the genus *Phytophthora*, such as the Irish Famine pathogen *Phytophthora infestans* and its tropical relative *Phytophthora palmivora*, cause multi-million-dollar losses yearly (Erwin and Ribeiro, 1996; Guest, 2007; Haverkort et al., 2009). Whether oomycetes display similar nuclear dynamics as filamentous fungi remains to be addressed. Pioneering work indirectly evidenced nuclear migration within germinating cysts from the diatom-infecting oomycete *Lagenisma coscinodisci* (Schnepf and Heinzmann, 1980). Similar to fungi, oomycete nuclei are known to distribute equally within coenocytic (aseptate) hyphae and this distribution is altered by the actin polymerization inhibitor latrunculin B (Ketelaar et al., 2012; Meijer et al., 2014; Fang and Tyler, 2017) or by silencing the *Phytophthora capsici* cell cycle regulator homologue *sda1* (Zhu et al., 2016).

We used time-lapse imaging to investigate nuclear dynamics of *P. palmivora* during cyst germination and subsequent root and leaf infection in addition to axenically cultured hyphae. For this purpose, we generated a versatile toolbox for efficient dual labelling of oomycete hyphae and organelles that enabled the rapid testing of various promoter-reporter constructs. We found that *P. palmivora* nuclei undergo coordinated bidirectional movements during plant infection and actively or passively move during mycelial growth to achieve a near-equal distribution of nuclei within hyphae. Active movement of individual nuclei frequently resulted in dramatic alterations of near-globular nuclei into extensively stretched shapes. Shape changes often co-occurred with a rerouting of nuclei toward nuclei-depleted hyphal sections. The centrosome-labelling Centrin2 protein occupied the trailing end of near-globular nuclei and the leading end of stretched nuclei. Furthermore, the microtubule polymerization inhibitor Benomyl decreased nuclear stretching and rerouting. Hence, our results unravel commonalities and differences in the dynamics of oomycete and fungal nuclei. Together, this constitutes the basis for new models of nuclear migration and hydrodynamic deformability that may ultimately offer alternative strategies to tackle oomycete diseases.

## Results

### Bi-directional nuclear movements occur during ***P. palmivora*** cyst germination

To investigate nuclear dynamics during *P. palmivora* cyst germination we generated the pTORKRm43GW Gateway vector which allows for dual labelling of hyphae and organelles **(Figures S1, S2, Table S1)** and transformed the *P. palmivora* isolate LILI (P16830). The vector contains a cassette for the Ham34-promoter-driven constitutive expression of a cytoplasmic tdTomato fluorescent reporter in addition to a nuclear-localized monomeric TFP1 (NLS:mTFP1) expressed under the *P. palmivora* ubiquitin-conjugating enzyme 2 (UBC2) native promoter **(Figures S3, S4)**. Among seven independent transformants, two showed distinct hyphal populations expressing either tdTomato or mTFP1 reporters, while five showed both cytoplasmic tdTomato (td) fluorescence and nuclear mTFP1 (NT) fluorescence. The latter were used to obtain the transgenic *P. palmivora* line ‘LILI-td-NT’ used in this study **Figure S3)**.

We then incubated encysted LILI-td-NT zoospores in a liquid compartment containing a *Nicotiana benthamiana* seedling **(Figure S5)** and monitored nuclear movements during the exploratory growth of cyst germ tubes (**Figure 1A-B, Video 1)**. Germ tubes emerged from mononucleate cysts and were initially devoid of a nucleus. After one hour when germ tubes were longer than 10 µm on average, the sole nucleus became stretched and moved from the cyst into the germ tube while the cyst remained filled with cytoplasm (**Figure 1A, Video 1)**. The nucleus divided in the germ tube but no leakage of mTFP1 fluorescence from the nucleus was detectable during division. The daughter nucleus N1 subsequently retracted to the cyst body, while the second daughter nucleus N2 moved forward within the germ tube at an average speed of 0.02 µm/s (**Figure 1A**, **Video 1)**. Upon further extension of the germ tube, the cyst-resident nucleus N1 re-entered the germ tube, still leaving cytoplasm in the cyst behind (**Figure 1A**, **Video 1)** and divided 3 hours after encystment into daughter nuclei N3 and N4 (**Figure 1B**, **Video 1)**. N3 and N4 moved away from each other, with nucleus N3 moving toward the cyst body and nucleus N4 moving forward within the germ tube. Collectively, these findings demonstrate that nuclei of germinating *P. palmivora* cysts divide in a closed mitosis and are then distributed evenly along the germ tube through concomitant, bi-directional movements.

**Figure 1.**
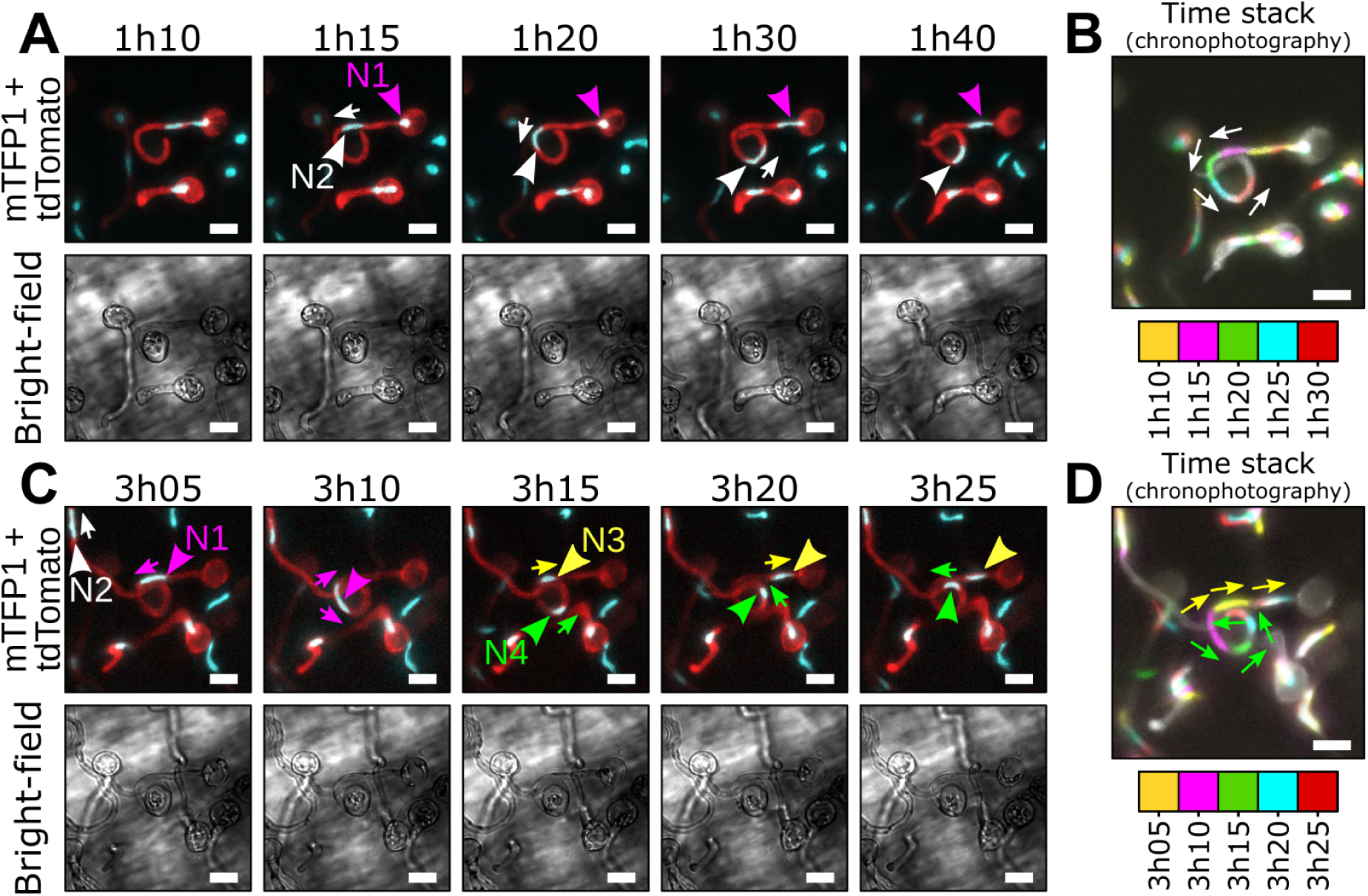
Repositioning of daughter nuclei follows division during *P. palmivora* cyst germination. **(A-D, Video 1)** Time-lapse imaging of germinating cysts from a transgenic *P. palmivora* strain expressing cytoplasmic tdTomato as well as nuclear-localized mTFP1 (LILI-td-NT) during exploratory growth at the surface of *N. benthamiana* roots. **(A)** Following first nuclear division, daughter nucleus N1 (magenta arrowhead) migrates toward the tip of the germ tube (yellow arrow), while the second nucleus N2 (white arrowhead) remains within the cyst body. **(B)** Chronophotographic view (time stack) of the previous sequence, showing the successive locations of nucleus N2. Time frames are color-encoded and overlaid. **(C)** Nuclear movements following N1 division. Daughter nuclei N3 (cyan arrowhead) and N4 (green arrowhead) move in opposite directions (yellow and white arrows, respectively) and distribute equally within the germ tube. **(D)** Chronophotographic view (time stack) of the sequence shown in **C**, showing opposite movement of nuclei N3 and N4. Scale bar is 10 µm.

### Appressorium differentiation alters nuclear dynamics in the germ tube

We then investigated nuclear movements during the onset of *N. benthamiana* root infection, using a similar set-up as previously described (**Figure 2**, **Figure S5, Video 2)**. Consistent with previous observations, the cyst-resident nucleus moved into the germ tube one hour after encystment, and reached the swollen tip of a differentiating appressorium. However, unlike cyst germination, we did not observe backward movement of a daughter nucleus towards the cyst after nuclear division. Instead, the cyst was progressively depleted from cytoplasm during appressorium differentiation (**Figure 2A**, **Video 2A)**.

**Figure 2.**
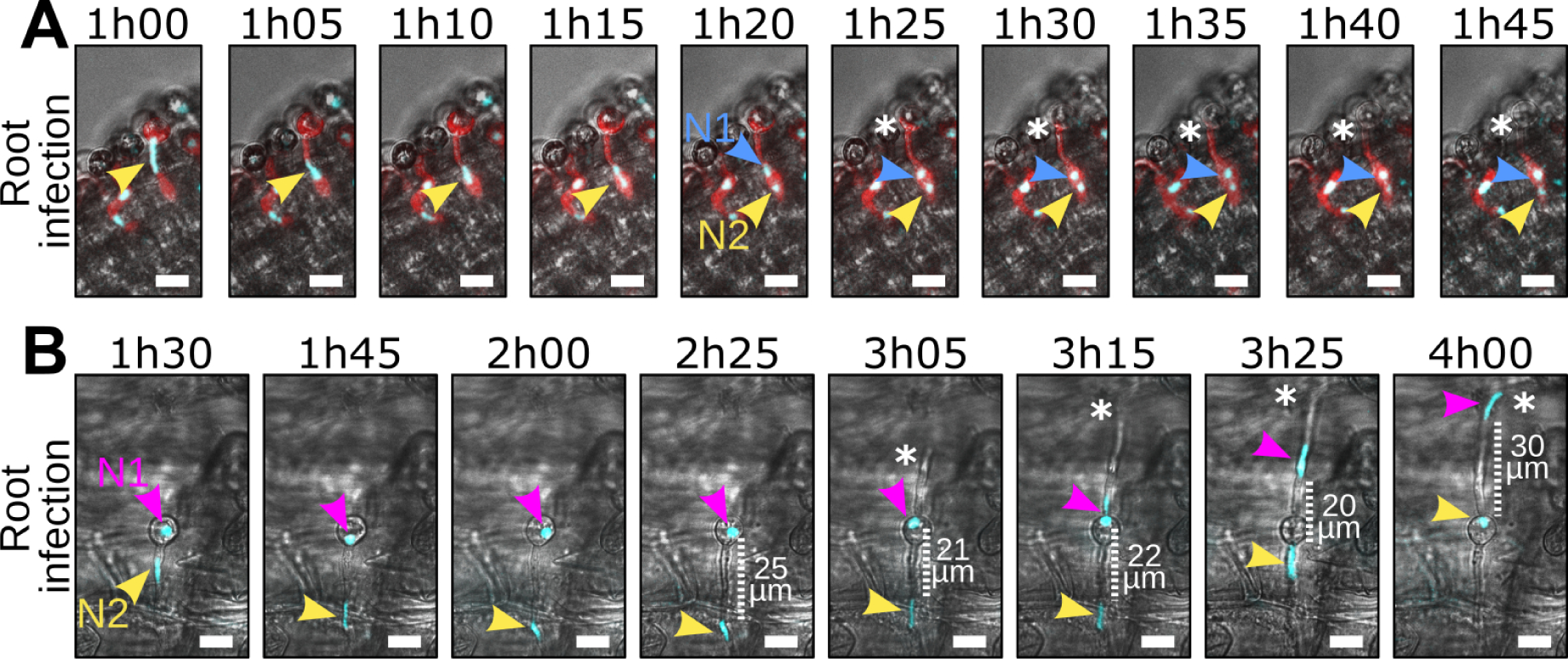
Appressorium differentiation alters *P. palmivora* nuclear dynamics in the germ tube. **(A-B, Video 2)** Time-lapse imaging of *N. benthamiana* root infection by a *P. palmivora* LILI-td-NT strain. **(A, Video 2A)** Representative sequence of a successful infection event. Arrowheads indicate nuclei. Asterisks indicate migration of the cytoplasm out of the cyst. **(B, Video 2B)** Representative sequence of an unsuccessful infection event, leading to the differentiation of a second germ tube (indicated by asterisks) opposite to the first one. Arrowheads indicate nuclei. Scale bar is 10 µm.

In a rare cyst germination event, appressorium differentiation did not occur (**Figure 2B**, **Video 2B)** and nuclear divisions and movements were similar to those during exploratory cyst germination, where nucleus N2 moved towards the germ tube tip and N1 resided in the cyst. Upon infection failure, a second germ tube germinated from the cyst opposite of the first after three hours. Nucleus N1 then proceeded from the cyst into the new germ tube, while nucleus N2 simultaneously retreated into the cyst. A distance of 20 to 30 µm was maintained between the two nuclei through the entire process (**Figure 2B**, **Video 2B)**. Taken together, these observations suggest that successful appressorium differentiation influences nuclear distribution dynamics, and that nuclei may be recalled and rerouted into additional infection attempts.

### Differential nuclear migration rates occur throughout the ***P. palmivora*** hyphal network both in planta and in axenic culture

During the biotrophic infection stage in living *N. benthamiana* leaves *P. palmivora* nuclei moved rapidly within intracellular hyphae (**Figure 3A-B**, **Video 3)**, but did not enter haustoria within plant cells (**Figure 3C-D, Video 4)**. By contrast to cyst germination, most nuclei adopted a round shape, likely due to the larger diameter of hyphae (up to 6 µm, N = 10) compared to germ tubes (2 µm, N = 10). Interestingly, independent hyphae showed differential nuclear migration rates, as evidenced by chronophotographic display of the time-lapse image series (**Figure 3B**). For instance, a continuous flow of nuclei was observed in the hyphal segment **b**, while no movement could be detected in the hyphal segment **a (Figure 3A**). Most nuclei were dragged passively by the bulk nuclear flow (P-type nuclei). Conversely, 10% of the nuclei, such as N1, stretched and maintained a near constant location over time within segment b despite a continuous flow of nuclei bypassing them (**Figure 3B**), which suggests that they are actively anchored (A-type nuclei). These findings demonstrate that different motion speeds occur throughout the hyphal network during biotrophy and that some nuclei may behave independently of the surrounding mass nuclear flow.

**Figure 3.**
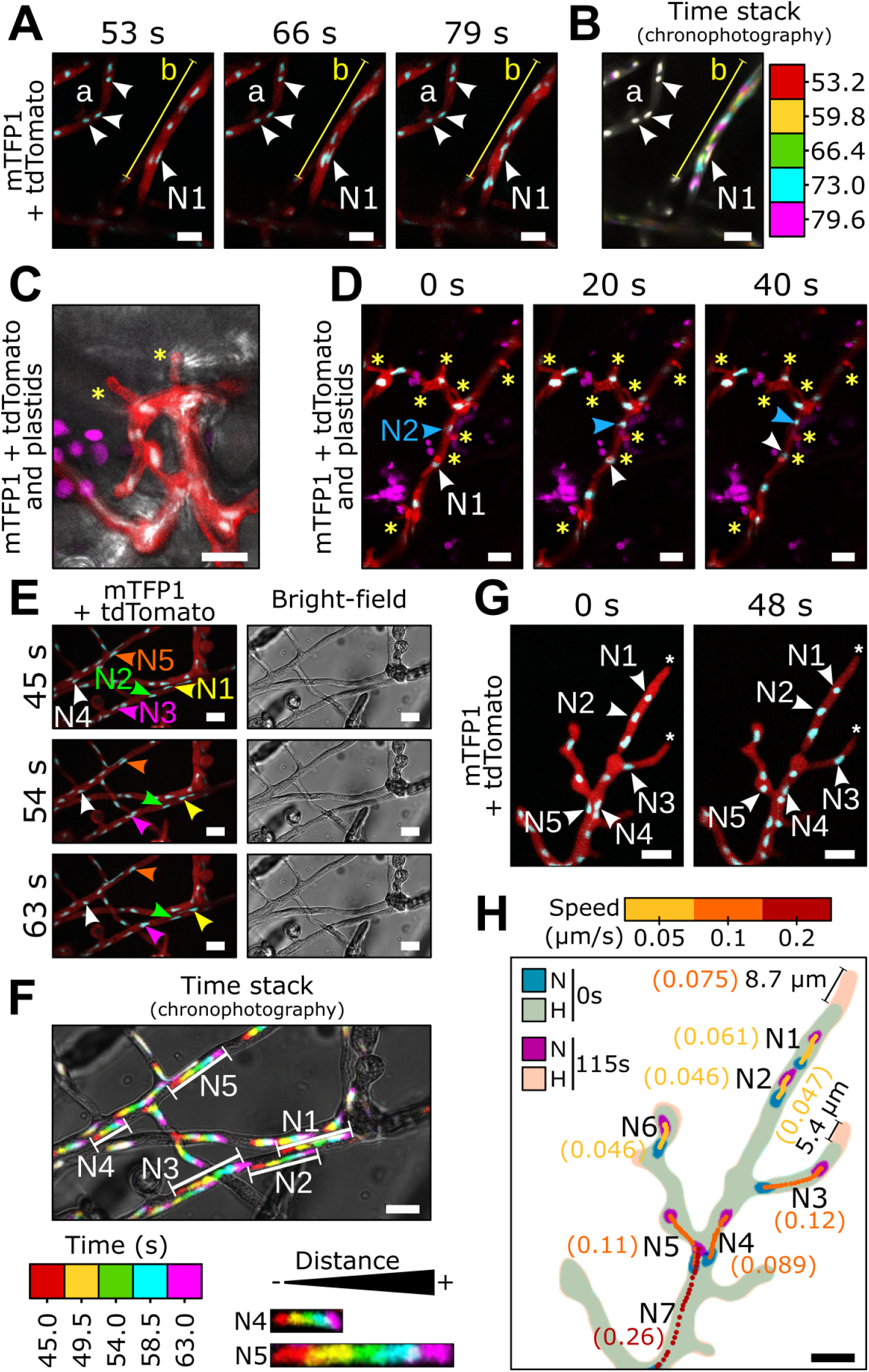
Differential nuclear migration rates occur throughout *P. palmivora* hyphal network. Time-lapse imaging of P. palmivora LILI-td-NT hyphae infecting a *N. benthamiana* leaf **(A-D, Videos 3-4)** or growing axenically on V8 medium **(E-H, Videos 5-6)**. **(A, Videos 3)** Representative sequence of nuclear movements within infectious *P. palmivora* hyphae. **(B, Video 3)** Chronophotographic display (time stack) of the sequence shown in **A**. The successive time frames are color-encoded and overlaid. **(C)** Representative sequence of *P. palmivora* haustoria (asterisks) on a *N. benthamiana* leaf. Arrowheads indicate nuclei. **(D, Video 4)** Representative sequence of nuclear movements within haustoriated *P. palmivora* hyphae. **(E, Video 5)** Representative sequence of nuclear migration on axenically-grown mycelium. **(F, Video 5)** Chronophotographic display of the sequence shown in **E**. **(G, Video 6)** Representative sequence of nuclear movements near hyphal tips. **(H)** Nuclear trajectories for seven nuclei at the hyphal tip. Average speed is indicated in brackets. Lines indicate hyphal segments. Arrowheads indicate individual nuclei. N: nuclei: H: hyphae. Scale bar is 10 µm.

Nuclear movements during saprotrophic growth on a V8 agar plate (**Figure 3E-H, Video 5**) were similar to those during plant infection and again differential nuclear motion speeds were observed within hyphal segments (**Figure 3E-F**). For instance, nuclei N1-N3 all moved with similar speed within the same hypha. By contrast, nucleus N5 moved faster than nuclei that moved passively with mass nuclear flow like N4. In addition, axenic hyphae allowed us to study hyphal tips (**Figure 3G-H, Video 6**), where mass nuclei movement slowed and nucleus N7 (average speed: 0.26 µm/s) moved faster than downstream nuclei N1-N6 (average speed ranging from 0.046 to 0.12 µm/s). Subapical zones were devoid of nuclei up to 9 µm (n = 10) from the hyphal tips (**Figure 3H**). Again, some nuclei migrate at speeds different from mass nuclear movement, likely due to active anchoring (A-type nuclei).

### A-type nuclei stretch and move independently of surrounding nuclear flow

To gain more insight on dynamics of A-type *P. palmivora* nuclei, we tracked nuclear stretching and other trajectories that differed from the mass nuclear movement and assessed their frequency within the hyphal network (**Figure 4**, **Video 5**). We identified several different behaviours of A-type nuclei. Frequently (28%), P-type nuclei transition into the A-type state and stop within the flow to either stay at a given position and stretch out sternwards (**Figure 4A**), suggesting they are anchored at one point, or moved against the flow (8.5%) (**Figure 4B**) and into branches (4%) (**Figure 4C**). In some rare cases (1.5%) (**Figure 4D**), A-type nuclei moving against the flow changed their direction of movement by tumbling, such that stretched part always pointed to the direction of the migration. With a frequency similar to the P- to A-type transition (28%), A-type nuclei reverted to passive movement (P-type) where they often shrank to adopt a round shape (**Figure 4E**). In rare cases (1.5%), such passive flow then also dragged these nuclei into side branches (**Figure 4F**). In summary, nuclei can undergo transitions between passive and active movements that allow them to position themselves within the hyphal network. Active movement correlates with changes in nuclear shape, where nuclei get remarkably stretched.

**Figure 4.**
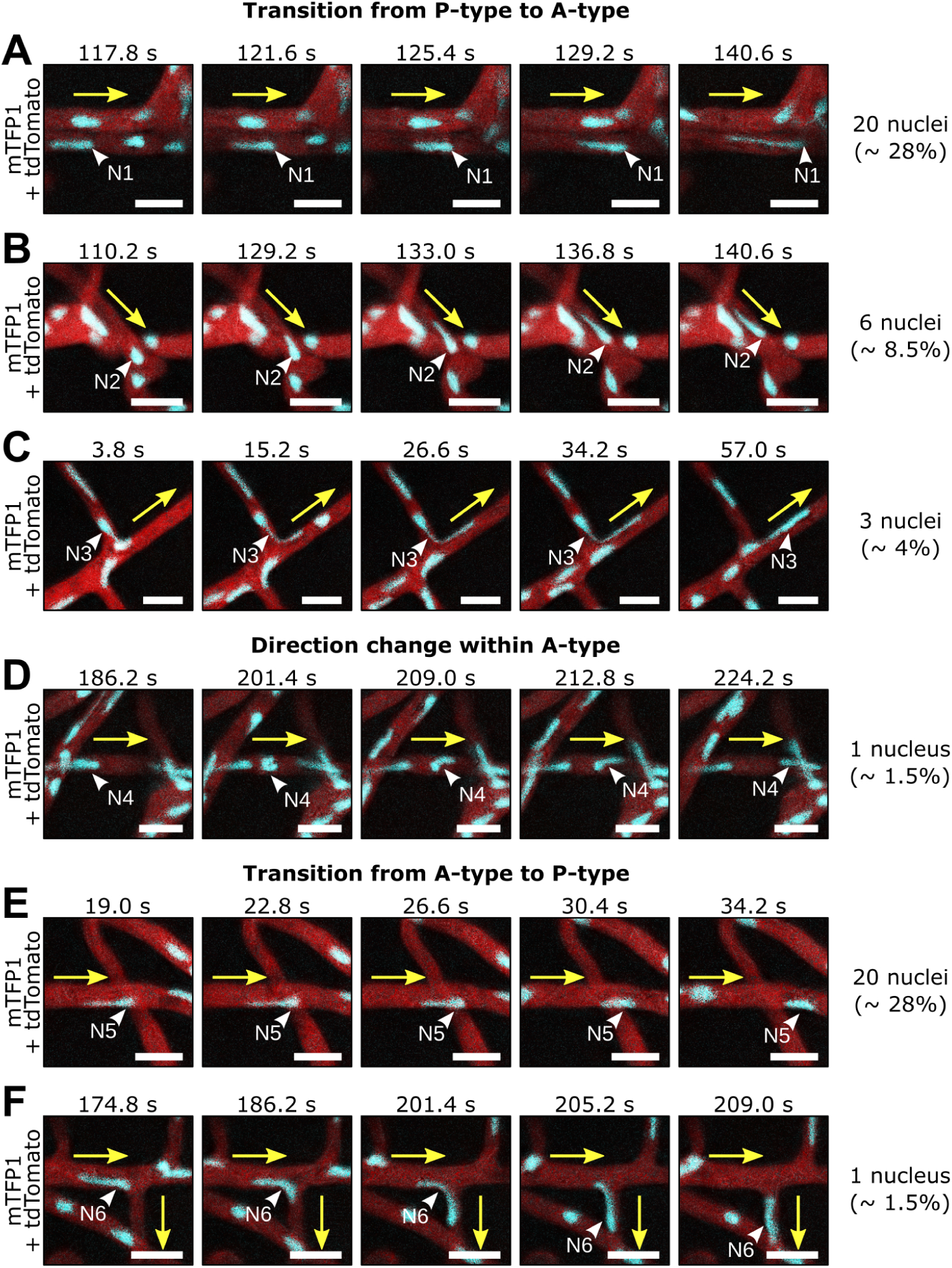
Individual *P. palmivora* nuclei stretch and move independently of surrounding nuclear flow. **(A-F, Video 6)** Time-lapse imaging of axenically-grown *P. palmivora* LILI-td-NT hyphae containing nuclei transitioning from passive (p-type) to active (a-type) movements **(A-D)** or conversely, from a-type to p-type **(E-F)**. **(A)** Nucleus slows down and stretches sternwards. **(B)** Nucleus shrinks and starts moving together with the surrounding nuclear flow. **(C)** Same as B, but nucleus enters a branch. **(D)** Nucleus stretches and moves opposite of the surrounding nuclear flow. **(E)** Same as **D**, but entering a branch. **(F)** Nucleus stretches and moves opposite of the surrounding nuclear flow, then tumbles and initiates a similar movement in the opposite direction. Yellow arrows indicate mass nuclear flow. White arrowheads indicate nuclei. Frequencies are given based on a total of 70 nuclei. Scale bar is 10 µm.

Nuclear shape flexibility is often associated with the absence of lamin-encoding genes (Xiang, 2018), but laminA has been identified in several *Phytophthora* species including *P. palmivora* (Kollmar, 2015; Ali et al., 2017). To test whether *P. palmivora* encodes a functional lamin A protein that controls nuclear shape, we generated transgenic strains expressing a mCitrine:laminA fusion under the control of either the constitutive Ham34 promoter or the native *laminA* promoter **(Figure S6)**. We found that expression of either construct resulted in a nucleoplasmic labelling with mCitrine. Notably, mCitrine:laminA overexpression triggered nuclear blebbing **(Figure S6A)** and arrested hyphal growth at an early stage, while expression under native promoter did not alter nuclear shape **(Figure S6B)** and hyphal growth was indiscernible from wild-type. Therefore, *P. palmivora* laminA may contribute on nuclear shape and integrity.

### Nuclear shape is altered upon acceleration and deceleration

To better understand how *P. palmivora* nuclei transition into A-type trajectories, we monitored the speed, shape and location of nuclei neighbouring individual nuclei at a hyphal branch point where the transition of P-to-A type may occur (**Figure 5**, **Video 7)**. We found that transitions between active and passive movement frequently associated with change in nuclear shape (**Figure 5B-D**) and facilitated a more equal nuclear distribution to fill empty areas (**Figure 5E-G**). For instance, nucleus N1 initially followed a P-type behaviour, then stretched and adopted an A-type behaviour, eventually moving backward into the branch point (**Figure 5B**) toward a 34 µm long hyphal segment devoid of nuclei (**Figure 5E**). Similarly, the stretched, A-type nucleus N2 moved backward into another branch point (**Figure 5C**), reaching a 35 µm long nuclear-depleted hyphal segment (**Figure 5F**). By contrast, the P-type nucleus N3 maintained a round shape while moving (**Figure 5D**) and an equal distance of 18 µm was maintained to the surrounding nuclei (**Figure 5G**).

**Figure 5.**
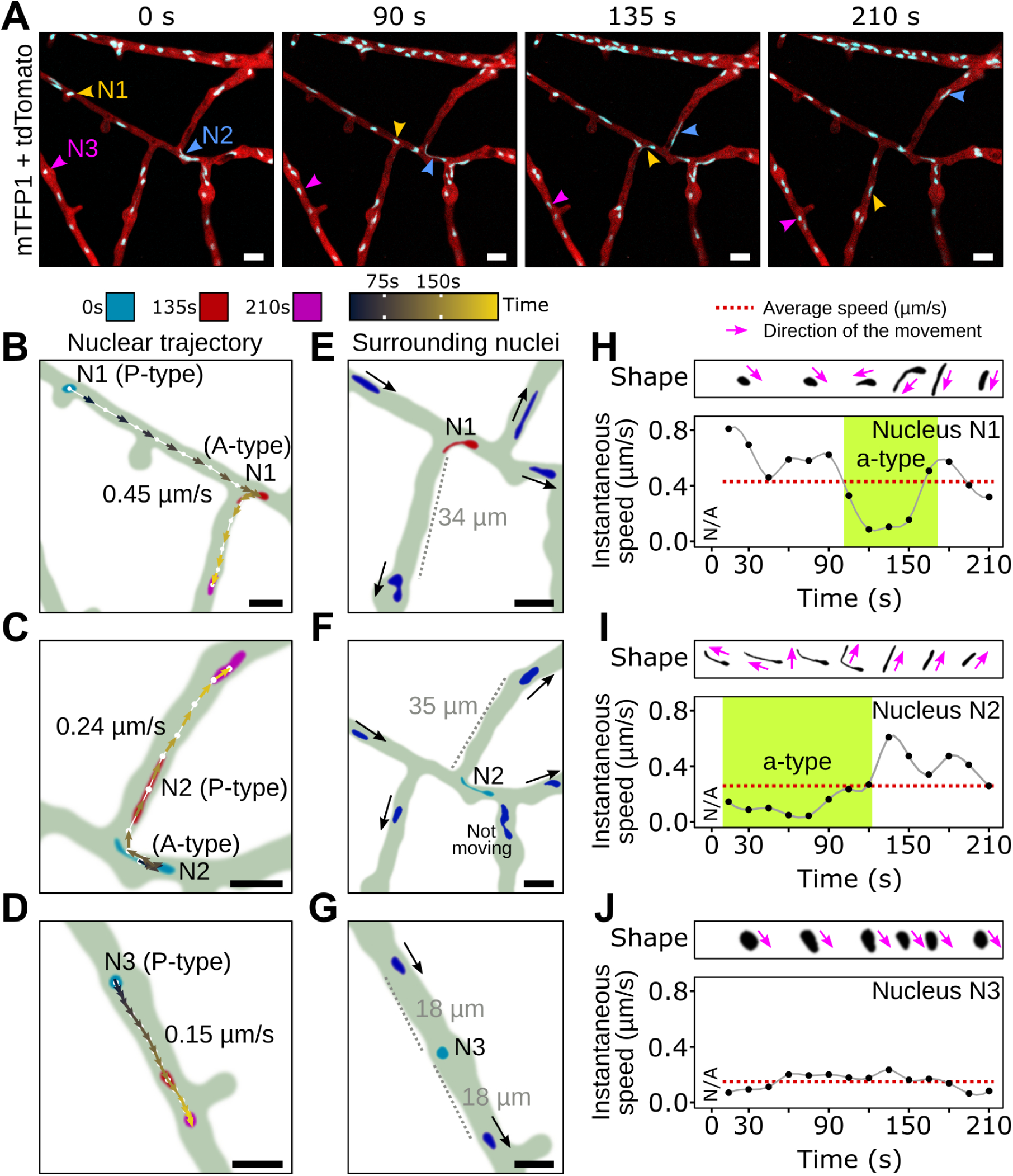
*P. palmivora* nuclear stretching correlates with rerouting toward nuclear-depleted hyphal segments. **(A, Video 7)** Representative sequence of nuclear movements at an hyphal branch point, high-lighting both a-type (nuclei N1 and N2) and p-type (nucleus N3) movements. Arrowheads indicate nuclei. **(B-J)** Analysis of nuclear speed and trajectories for nuclei N1 to N3. **(B, E, H)** Maps of nuclear content in the vicinity of nuclei N1 **(B)**, N2 **(E)** and N3 **(H)**. Black arrows indicate nuclear movement. **(C, F, I)** Nuclear trajectories of N1 **(C)**, N2 **(F)** and N3 **(I)**. Locations of nuclei N1 to N3 at first (0 s), intermediate (135 s) and last (210 s) time points are shown in blue, red and magenta, respectively. Nuclei centroids are represented as dots. Arrows indicate instantaneous speed and direction of the movement. Average nucleus speed is indicated on the graph. **(D, G, J)** Variation of instantaneous speed with time for nuclei N1 **(D)**, N2 **(G)** and N3 **(J)**. Average speed in shown as a red dot line. Representative pictures of nucleus shape over time are shown above the graphs with arrows indicating direction of the movement. Scale bar is 10 µm.

We then measured instantaneous speed and direction of nuclei undergoing significant shape changes (**Figure 5H-J**). We found that the onset of shape change in A-type nuclei correlated with changes in their instantaneous speed and that shape was affected by the direction of movement. For instance, nucleus N1 stretched while undergoing a ten-fold speed decrease from 0.8 µm/s down to 0.08 µm/s and reverted to a round shape after speed increased back to 0.4 µm/s (**Figure 5H**). Similarly, stretched nucleus N2 had an initial speed of 0.1 µm/s and later became more round after speed increased (**Figure 5I**). By contrast, the P-type nucleus N3 that maintained a steady speed did not change its shape (**Figure 5J**). Taken together, these results suggest that trajectories of A-type nuclei lead to their reallocation of nuclei into nuclear-depleted hypha sections. The co-occurrence of nuclear stretching during these reallocations suggests that forces act most prominently at a single nuclear envelope pole.

### A-type nuclear movement is impaired by the microtubule-polymerization inhibitor Benomyl

Since microtubules are required for nuclear movements in fungi (Morris et al., 1995; Xiang, 2018), we assessed the effect of the microtubule-polymerization inhibitor Benomyl on *P. palmivora* nuclei (**Figure 6**, **Video 8**). Benomyl restricted *P. palmivora* growth in a dose-dependent manner. While 50 mg/L Benomyl arrested hyphal growth on V8 agar plate and resulted in abnormally-shaped hyphae (**Figure S7**), 10 mg/L Benomyl barely reduced *P. palmivora* growth (**Figure 6A-B**) and did not alter hyphal shape (**Figure 6C-D**). Thus, 10 mg/L benomyl can be used to assess *P. palmivora* nuclear movements independent of changes in overall growth rate and hyphal anatomy. A near equal distribution of nuclei was maintained along hyphae, suggesting that such a concentration did not have a dramatic effect on *P. palmivora* physiology. Time-lapse imaging showed nuclear movements in Benomyl-treated hyphae (**Figure 6C-E**, **Video 8B)**. However, only 14% of nuclei displayed A-type behaviour upon Benomyl treatment, compared to 42% in the control (N = 23, 400 nuclei) (**Figure 6E**, **Video 8A)**. Instead, most nuclei (86%) were round and followed P-type trajectories, compared to 58% in the control (**Figure 6E**). Taken together, these findings suggest that microtubules are required for A-type nuclear movements in *P. palmivora*.

**Figure 6.**
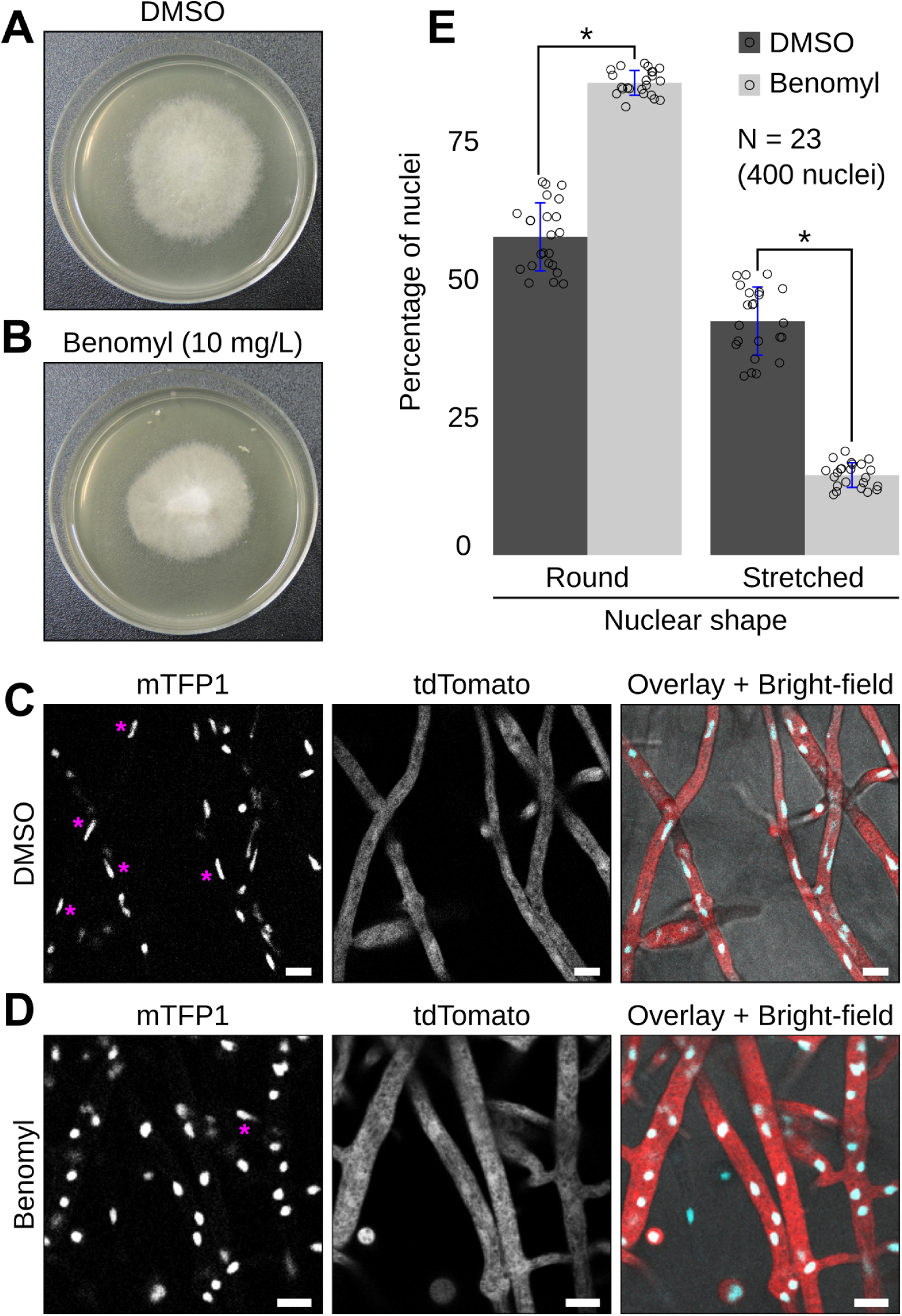
Antimicrotubule drug benomyl impairs *P. palmivora* A-type nuclear movements. **(A-B)** Growth habit of transgenic *P. palmivora* LILI-td-NT mycelium growing on V8 agar plates without (DMSO control) **(A)** or with **(B)** with 10 mg/L Benomyl. (C-D) Representative pictures of *P. palmivora* LILI-td-NT hyphae and nuclei in absence **(C, Video 8A)** or presence **(D, Video 8B)** of 10 mg/L Benomyl. Scale bar is 10 µm. **(E)** Quantification of round (P-type) and stretched (A-type) nuclei within *P. palmivora* LILI-td-NT hyphae (N = 23, 400 nuclei). Statistical significance was assessed using Wilcoxon test for paired samples (*P* < 0.05).

### The centrosome-associated protein Centrin2 localizes at the stretched extremities of nuclei

Fungal research has demonstrated that cell cortex-anchored dynein pulls the astral microtubules nucleated from the SPBs to move nuclei (Morris et al., 1995; Xiang, 2018). Centrosomes and centrioles which could serve as SPB equivalent in oomycetes have been reported for several *Phytophthora* species (Heath, 1974). We therefore labelled *P. palmivora* centrosomes using a fluorescently-tagged centriolar lumen protein Centrin2 (CETN2). To that end, we generated a transgenic *P. palmivora* ‘LILI-NT-Ce’ strain by replacing the cytoplasmic tdTomato cassette from the previous dual reporter construct with a mCitrine fluorescent reporter fused N-terminally to *P. palmivora* CETN2 coding sequence (Ce) and monitored fluorescence over time at branching hyphae (**Figure 7**, **Video 9)**. CETN2 is composed of a nuclear localization signal followed by a Ca2+-binding protein/EF-hand superfamily protein domain (PTZ00183) (**Figure 7A**). Expression of the CETN2/nuclei dual reporter led to labelling of nuclei with two adjacent punctate structures at the nuclear periphery (**Figure 7B**) likely corresponding to the two centrioles of the *P. palmivora* centrosome (**Figure 7C**). Similar labelling was obtained with the constitutive Ham34 promoter as well as the native *P. palmivora* CETN2 promoter (**Figure S8**). In addition, up to 15% of observed nuclei within a given region showed two sets of punctate structures located opposite of each other, presumably as a result of centrosome duplication prior to nuclear division (**Figure 7D**, **Figure S9**).

**Figure 7.**
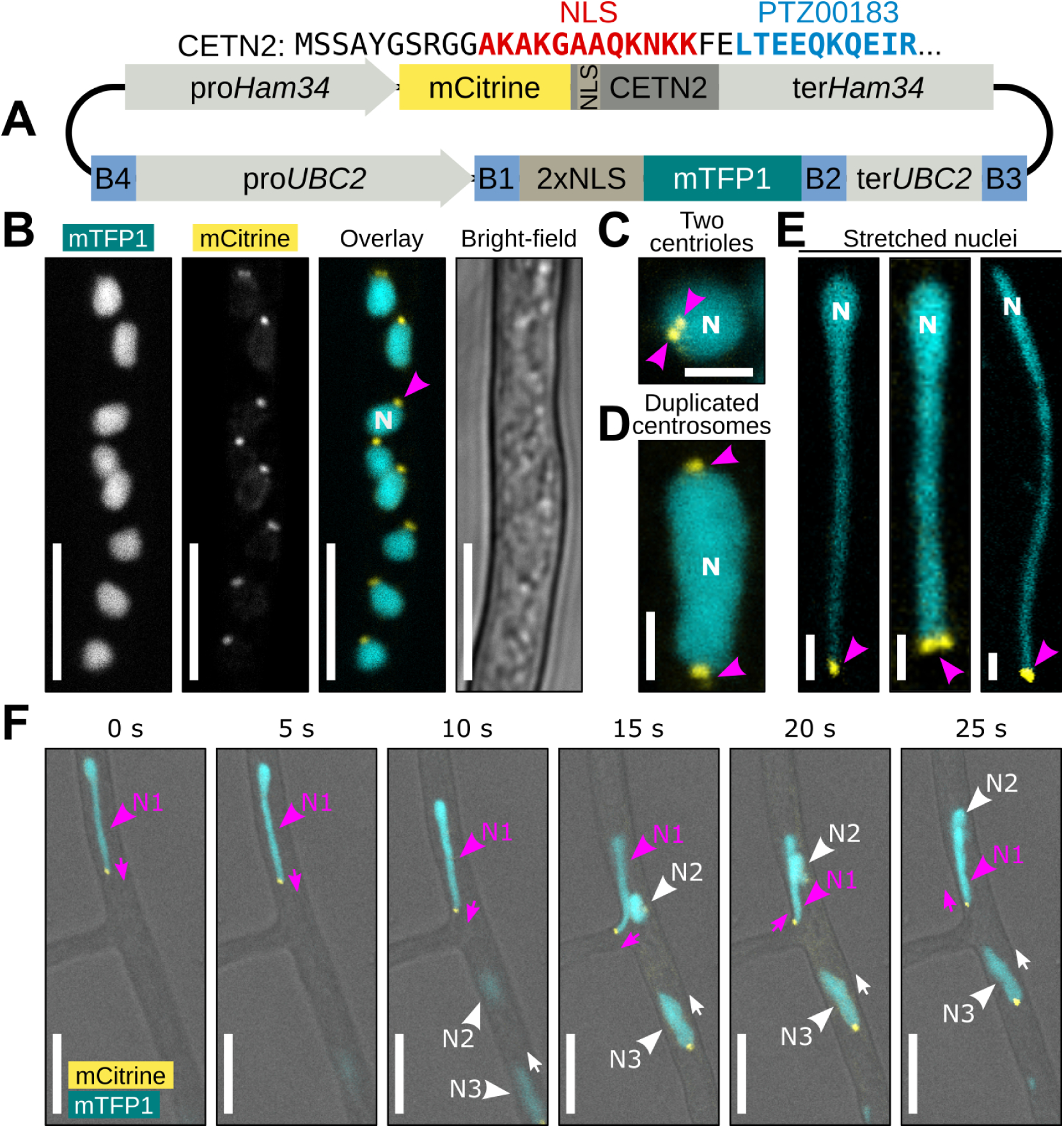
Centrin2 localizes to *P. palmivora* nuclear stretches. **(A)** Schematic view of the construct used for dual labelling of nuclei and centrosomes in *P. palmivora* strain LILI-NT-Ce. **(B, Video 9)** Representative picture of mCitrine-labelled Centrin2 (CETN2) within *P. palmivora* LILI-NT-Ce hyphae. **(C)** CETN2 localizes in two adjacents dots at the periphery of the nucleus. **(D)** Centrosome duplication in dividing nuclei. **(E)** Representative pictures of stretched *P. palmivora* nuclei (up to 30 µm long), with CETN2 localizing at the tip of the stretched areas. **(F)** Time-lapse imaging of nuclei movements at hyphal branch point. Nucleus N1 (magenta) follows a a-type trajectory backward to the branch point, while nuclei N2 and N3 follow a p-type trajectory. CETN2 localizes at the tip of the stretched part of nucleus N1, while CETN2 localizes at the back of nuclei N2 and N3. Scale bar is 2 µm **(C-E)** or 10 µm **(B, F)**.

We then specifically looked for elongated nuclei within *P. palmivora* hyphae and found several instances of *P. palmivora* nuclei stretching up to 30 µm in length. In all cases, mCitrine:CETN2-labelled punctate structures occurred at the very tip of the nuclear stretch (**Figure 7E**). We then investigated the relationships between nuclear movements and positioning of mCitrine:CETN2-labelled punctate structures. We found that migrating P-type nuclei had their punctate structures oriented sternwards, that is opposite of the direction of movement (nuclei N2 and N3) (**Figure 7F**, **Video 9)**. By contrast, stretched nuclei moving against the mass nuclear flow had their punctate structures oriented in front (nucleus N1) (**Figure 7F**). Our results suggest that *P. palmivora* nuclei are highly deformable and mCitrine:CETN2-labelled punctate structures may be involved in microtubule-aided nuclear anchorage within hyphae and their movement against cytoplasmic flow.

## Discussion

We studied the broad-host-range plant pathogen *P. palmivora* as a system to elucidate oomycete nuclear dynamics. By combining live imaging and a dual reporter expression system, we provide evidence for both active and passive nuclear movements within multi-nucleate, coenocytic hyphae. Different nuclear migration rates occur within the same hyphal segments and nuclei move at a reduced rate in subapical regions comparing to other hyphal segments, suggesting the overall nuclear movement is influenced by cytoplasmic flow toward the mycelium periphery. Consistent with this hypothesis, nuclear migration in the fungus *Neurospora crassa* involves cytoplasmic flow (Lew, 2005; Ramos-García et al., 2009) and can be reversed by application of osmotic gradients across the colony (Roper et al., 2013). However, by contrast to observations made on *Aspergillus nidulans* (Fischer, 1999), we did not observe a saltatory movement of nuclei, *i.e.* periodic inversion of the mass nuclear flow.

We describe the active, long-distance rerouting of individual nuclei toward nuclear-depleted areas and our data suggest that such A-type movements contribute to the near equal spacing of nuclei within germ tubes, infectious hyphae and axenic mycelium (**Figure 8**). We found that A-type movements are impaired by Benomyl, suggesting they require microtubules. Consistent with our observations, a proper nuclear distribution within *A. nidulans* and *N. crassa* hyphae requires microtubules (Morris, 1975; Oakley and Morris, 1980) and an additional role of cytoplasmic dynein has been reported (Plamann et al., 1994; Xiang et al., 1994). Future work will investigate the role of *P. palmivora* dynein in nuclear movement.

**Figure 8.**
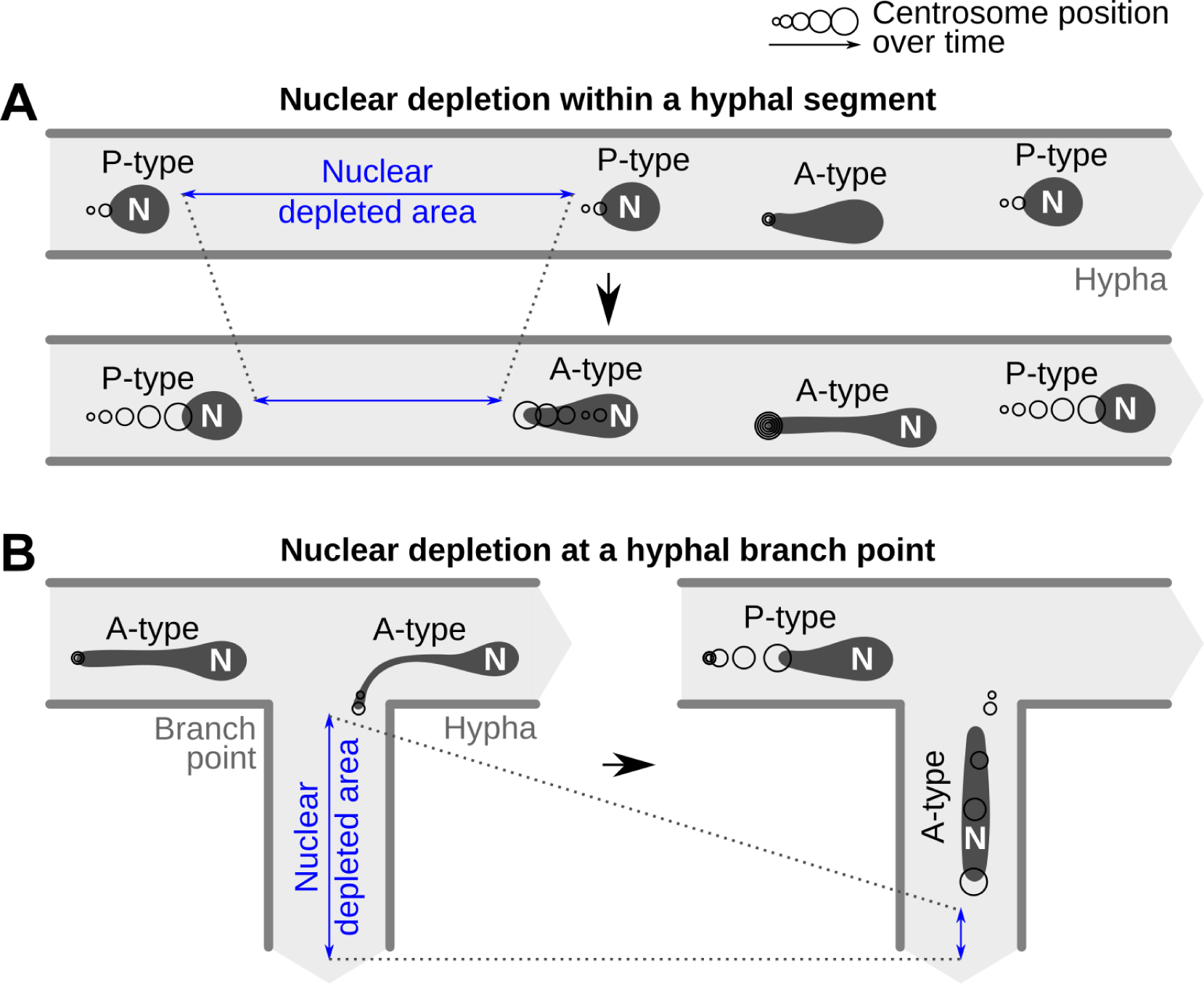
Proposed behaviour change from passive (P-type) to active (A-type) movements within *P. palmivora* hyphae. **(A)** Decrease in nuclear density within an hyphal segment (blue) results in down-stream nuclei to anchor until populated by P-type nuclei or initiate retrograde movement to fill it directly. Anchorage is mediated by the centrosome and results in nuclear stretching. **(B)** Decrease in nuclear density in a branching hypha (blue) results in nuclei to anchor at the branch point and initiate active movement to enter the branch.

Several reports suggest that a local variation in tubulin concentration may affect micro-tubule polymerization in different model systems. For instance, a local increase in tubulin concentration in neurons may favour nucleation of non-centrosomal microtubule bundles (Hernández-Vega et al., 2017). Besides, the onset of mitosis in the fungus *A. nidulans* triggers a rapid influx of tubulin into the nucleoplasm (Ovechkina et al., 2003) and local accumulation of tubulin monomers in the nascent spindle region of mitotic nuclei has been reported from *Caenorhabditis elegans* embryos (Hayashi et al., 2012). The tempting speculation that local changes in tubulin concentration favour transition from P- to A-type nuclear trajectories in *P. palmivora* hyphae will require future experimentation.

Equidistant nuclear distribution mechanisms and nuclear migration are likely to occur in other oomycete species, including those not belonging to the family *Peronosporaceae*. For instance, indirect evidence suggest that nuclear migration occurs in germinating cysts of *Lagenisma coscinodisci*, a diatom parasite from an early diverging Saprolegniomycetes lineage (Thines et al., 2015). In this species, a near-globular nucleus migrates from the cyst to the germ tube and concomitantly stretch, reaching a length of up to 10 µm, presumably due to the narrowness of the germ tube (Schnepf and Heinzmann, 1980). Therefore, *P. palmivora* nuclear dynamics may be similar to other oomycetes.

We showed a correlation between active movement of *P. palmivora* nuclei and nuclear stretching. A-type nuclei were stretched up to 30 µm in length upon speed change, irrespective of the diameter of the surrounding hyphae. They revert back to round shape within seconds when returning to P-type trajectory. Nuclear deformability has also been reported in fungi. For instance, during rice infection by *Magnaporthe oryzae*, migration of the appressorial nucleus within the narrow penetration peg causes nuclear constriction up to 13 µm (Jenkinson et al., 2017). While extreme constrictions of fungal nuclei were triggered by the narrowness of the surrounding hyphal structures (Jenkinson et al., 2017), *P. palmivora* nuclei also deformed in hyphae with diameters much larger than their nuclei. Deformability has been associated with a specific nuclear lamina composition and chromatin organization (Davidson and Lammerding, 2014). Variation in lamin gene expression regulates nuclear stiffness and its deformability in a variety of metazoan cells, including hematopoietic and immune cells (Bone and Starr, 2016). Lamin overexpression increases nuclear stiffness and reduces deformability (Bone and Starr, 2016), while loss of lamin gene expression in metazoans resulted in increased nuclear deformability but also in increased fragility (Rowat et al., 2013; Davidson and Lammerding, 2014). Moreover, defects in lamina assembly are responsible for increased cell death and cause a large array of life-threatening laminopathies (Davidson and Lammerding, 2014) and lamin null mutants show decreased nuclear stiffness. Fungal nuclei are devoid of lamina (Cardenas et al., 1990) and lack lamin-encoding genes (Kollmar, 2015), presumably explaining their deformability. In contrast to fungi, lamin-encoding genes are present in several *Phytophthora* species including *P. infestans*, *P. ramorum* and *P. palmivora* (Kollmar, 2015; Ali et al., 2017). Furthermore, we showed that overexpression of the *P. palmivora* laminA gene alters the nuclear periphery and affects nuclear shape formation in *Phytophthora*.

The centrosome of stretched A-type nuclei always occupies the leading tip suggesting that nuclear stretching is a result of direct or indirect forces exerted on the centrosome. In agreement with this hypothesis, we observed nuclear tumbling, indicative that A-type movement is polarized. Finally, the astern orientation of centrosomes in P-type nuclei suggests that passive movement may be a transient state that prepares for possible conversions to A-type for nuclear rerouting. Taken together, *P. palmivora* is an accessible model organism to study the relationships between nuclear movements, deformability and rerouting.

Models of lipid bilayer vesicles flowing through circular tubes (Barakat and Shaqfeh, 2018; Kaoui et al., 2011; Aouane et al., 2014) do not predict shapes like those observed in A-type nuclei but primarily reflect red blood cell microcirculation in capillaries and liposome flow through microfluidic compartments. These models therefore do not take into account nuclear-specific parameters such as interactions with the cytoskeleton (Dickinson et al., 2015). Besides, simple lipid bilayers may not reflect the biophysical properties of nuclear envelope. For instance, biomechanical models of nuclear shape changes during micropipette aspiration showed that the overall response of an isolated nucleus is highly sensitive to the apparent stiffness of the nuclear lamina (Vaziri and Mofrad, 2007). We speculate that dramatic stretching minimizes the constraints exerted on nuclei allowing them to migrate greater distances with limited energy cost. Future work should aim at developing models for hydrodynamic shape adaptations of nuclei to sustain long-distance, flow-independent movements within branched filamentous hyphae.

We documented bi-directional movements of nuclei during cyst germination which were altered by appressorium differentiation. Failure to differentiate an appressorium triggered rerouting of nuclei into additional infection attempts. This suggests that *P. palmivora* nuclear dynamics and developmental transitions are coordinated. Similarly, during rice infection by the fungus *M. oryzae*, one post-mitotic nucleus migrates into the developing appressorium, while other nuclei is degraded after entering the conidial cell (Vaneault-Fourrey et al., 2006). Later on, the appressorial nucleus migrates down into the primary hypha. Such migration requires a functional transketolase TKL1 acting as a metabolic check-point. Indeed, loss of transketolase function results in depletion of ATP content and slowing down of TOR signaling pathway (Fernandez et al., 2014). Whether an expressed transketolase ortholog of *TKL1* in *P. palmivora* (PLTG_04838) (Evangelisti et al., 2017) is also in-volved in early infection processes remains to be addressed.

In conclusion, *P. palmivora* is an accessible system which allowed us to highlight commonalities as well as differences in nuclear dynamics between oomycetes and previously studied fungi. Our work sheds new light on nuclear deformability in filamentous microorganisms and uncovers the role of nuclear rerouting to dynamically maintain equal distribution within coenocytic hyphae as well as during plant infection. These findings provide a foundation for hydrodynamic modelling of nuclear shape adaptations in branched hyphal networks and will contribute to develop robust, hard-to-evade targets for durable crop protection against oomycete diseases.

## Materials and methods

### Phytophthora strains and growth conditions

*P. palmivora* Butler isolate LILI (accession P16830) was initially isolated from oil palm in Colombia (Torres et al., 2010) and maintained in the *P. palmivora* collection at the Sainsbury Laboratory (Cambridge, UK). *P. palmivora* strains were maintained on Petri plates of V8 agar (1.5% agar) at 25°C. For maintenance of transformed strains, geneticin (G418) was added to the medium at a concentration of 100 mg/L. Mycelium was grown for 5 days in the dark followed by 2 days under constant light conditions. For the latter, plates were left unsealed to remove excess humidity. For production of zoospores, 7-day-old plates were incubated at 4°C for 30 min. Plates were then flooded with 5 mL sterile water and incubated at room temperature for another 30 min. *N. benthamiana* growth conditions were described previously (Evangelisti et al., 2017).

### Plasmid construction

An attR3/attR4 MultiSite Gateway cassette was PCR-amplified from pK7m34GW (Karimi et al., 2005) using GoTaq DNA Polymerase (Promega UK, Southampton, UK) with primers NdeI-R3 and NdeI-R4 **(Table S2)**, flanking each side of the cassette with NdeI restriction sites. The MultiSite Gateway cassette was then ligated into pGEM-T Easy Vector (Promega UK, Southampton, UK) and the insert sequence was confirmed by sequencing (Source BioScience, Nottingham, UK). Thereafter the cassette was ligated into pTOR vector (Judelson and Michelmore, 1991) using the NdeI restriction sites replacing the vector’s Ham34 promoter, multiple cloning site and Ham34 terminator, resulting in plasmid pTORKm43GW. The orientation of the attR3/attR4 cassette was selected so promoters inserted using the Gateway cassette will not be in close proximity to the Hsp70 promoter driving antibiotic selection marker. pTOR vectors carrying a cassette for Ham34 promoter-driven expression of mTFP1 (cyan) (Ai et al., 2006), mWasabi (green) (Ai et al., 2008), mCitrine (yellow) (Griesbeck et al., 2001) or tdTomato (red) (Shaner et al., 2004) were derived from KpnI-linearized pTORKm43GW. Briefly, Ham34 promoter, Ham34 terminator and the reporters were amplified individually using primer pairs listed in **Table S2**. The final cassettes were assembled by overlap extension PCR and used for In-Fusion cloning (Clontech, Palo Alto, USA) into linearized pTOR-Gateway vectors. An outline of the pTOR-Gateway backbone is shown in **Figure S1**.

### Genomic DNA extraction

Genomic DNA was extracted from axenically-grown *P. palmivora* mycelium using a protocol modified from Möller et al. (1992). Briefly, samples were incubated in a 1:1 mix of STES buffer (50 mM Tris-HCl pH 8.0, 10 mM EDTA, 150 mM NaCl, 2%(v/v) SDS) and Trisbuffered phenol solution (pH 8.0) for 30 min at 65°C. After phenol-chloroform extraction, nucleic acids were treated with RNase T1 (Life Technologies Ltd., Paisley, UK) for 30 min at 37°C. RNase was removed by phenol-chloroform extraction prior to isopropanol precipitation of genomic DNA.

### Cloning of Phytophthora promoters and terminators

Primers for *P. palmivora* promoters and terminators used in this study were derived from publicly available genomic resources (Ali et al., 2017) **(Table S2)**. Constructs were generated using two-step Gateway PCR (Invitrogen, Carlsbad, USA) and subsequently cloned into pDONR221 entry vectors carrying attP4-attP1R or attP2R-attP3 Gateway cassette, respectively. Sequences were confirmed by sequencing (Source BioScience, Nottingham, UK). The ubiquitin-conjugating enzyme 2 (UBC2) promoter was defined as 1500 bp upstream of start codon. Terminator consisted of 500 bp downstream of UBC2 stop codon. LaminA and Centrin2 (CETN2) promoters were defined as 1000 bp ustream of start codon. The nuclear reporter used in this study was obtained by fusing a tandem repeat of *Phytophthora sojae* bZIP1 nuclear localization signal (Fang and Tyler, 2017) to the 5’ end of mTFP1. The centrosome reporter was obtained by fusing mCitrine to the 5’ end of *P. palmivora* CETN2.

### Generation of transgenic *P. palmivora*

Transgenic *P. palmivora* were obtained by zoospore electro-transformation using the method from Huitema et al. (2011) with the following modifications: for electroporation, 680 µL of high concentration (>10^6^ zoospores/ml), high mobility zoospore suspension was mixed with 80 µL of 10*×* modified Petri’s solution and 40 µL (20-80 µg) of plasmid DNA. Electroporation settings were as follows: voltage 500 V, capacitance 50 µF, resistance 800 ohms. After electroporation, zoospore suspensions were diluted to 6 mL with clarified V8 medium and incubated at 25°C for 6 h on a rocking shaker. The encysted zoospore suspension was plated on a 15 cm diameter plate with selective medium containing 100 mg/L geneticin. Transformants were transferred to fresh selective plates up to 10 days after transformation.

### Quantitative reverse transcription-polymerase chain reaction (qRT-PCR) analyses

Total RNA was extracted from axenically-grown *P. palmivora* mycelium containing sporangia (sample MZ) and *N. benthamiana* roots harvested at 3, 6, 18, 24, 30 and 48 h after inoculation (hai) with *P. palmivora* ARI-td zoospores (Le Fevre et al., 2016) using the RNeasy Plant Mini Kit (Qiagen, Germantown, USA). One microgram was reverse transcribed to generate first-strand complementary DNA (cDNA), using the Bio-Rad IScript cDNA Synthesis Kit according to the manufacturer’s instructions (Bio-Rad, Hercules, USA). RNA quality was assessed by electrophoresis on agarose gel. Conditions for quantitative PCR were described previously (Evangelisti et al., 2017).

### Confocal microscopy

Confocal laser scanning microscopy images were acquired with a Leica SP8 laser-scanning confocal microscope equipped with a Leica HC FLUOTAR 25*×* 0.95 numerical aperture (NA) objective (Leica, Wetzlar, Germany). A white-light laser was used for excitation at 477 nm for mTFP1 visualisation, 488 nm for mWasabi visualisation, at 514 nm for mCitrine visualisation and at 543 nm for the visualisation of tdTomato. Fluorescence acquisition was done sequentially. For time-lapse imaging of infected N. benthamiana roots, seedlings were mounted between a slide and a coverslip in sterile water containing the zoospore suspension. Coverslip was sealed to the slide using a modified Valap sealing (Chazotte, 2011) composed of a 4:1 mix of paraffin (Sigma-Aldrich, UK) and lanolin (Sigma-Aldrich, UK) to prevent dehydration **(Figure S5)**. Time-lapse imaging of axenically-grown mycelium was carried out on plates flooded with 10 mL sterile water. Pictures were analysed with the ImageJ software (http://imagej.nih.gov/ij/) using the plugin Bio-Formats (https://imagej.net/Bio-Formats). Signal-to-noise ratio was optimized uniformly on time-lapse images series by adjusting minimum and maximum intensity levels. Then, Z-stacks were overlaid using maximum intensity projection and saved in false colors. Videos were generated from overlaid images with ffmeg (https://ffmpeg.org/).

### Measurements of nuclear speed and shape

Nuclear position was defined as the coordinates of its centroid, obtained from the ImageJ software. Instantaneous speed was plotted using the R software (https://www.r-project.org/) and ggplot2 package (https://ggplot2.tidyverse.org/). Nuclear shape was defined as the ratio (R_F_) between Feret’s maximum diameter (maximum caliper) and Feret’s minimum diameter (minimum caliper). Nuclei were considered round when R_F_ < 2, while larger R_F_ values indicated stretched nuclei.

### Online supplemental material

**Figure S1** outlines the pTOR-Gateway backbone. **Figure S2** shows growth habit of transgenic *P. palmivora* strains carrying empty pTOR-Gateway vectors during *N. benthamiana* leaf infection. **Figure S3** illustrates the variability of primary *P. palmivora* transformants carrying a dual hyphae/nuclei reporter. **Figure S4** shows transcripts levels corresponding to the *P. palmivora* UBC2 gene on mycelium with sporangia (MZ) in addition to plant infection. **Figure S5** shows the experimental set-up used for time-lapse imaging of *N. benthamiana* root infection. **Figure S6** shows alterations of nuclear shape in transgenic *P. palmivora* strains expressing a laminA-reporter either constitutively or under its native promoter. **Figure S7** shows the effect of Benomyl on growth and development of *P. palmivora* hyphae. **Figure S8** compares subcellular localization of a Centrin2 reporter expressed either constitutively or under its native promoter. **Figure S9** illustrates the frequency of *P. palmivora* nuclei with duplicated centrosomes within hyphae.

### Video legends

**Video 1.** Time-lapse confocal microscopy of germinating *P. palmivora* cysts from a transgenic strain expressing a cytoplasmic tdTomato and a nuclear-localized mTFP1 (LILI-td-NT) during exploratory growth at the surface of *N. benthamiana* roots. Z-stacks were collected every 5 min. Still images corresponding to this video are displayed in **Figure 1**. Scale bar is 10\µm.

**Video 2.** Time-lapse confocal microscopy of *P. palmivora* LILI-td-NT infection structures during *N. benthamiana* root infection. The video is composed of two movies representing successful **(Video 2A)** and unsuccessful **(Video 2B)** infection events. Z-stacks were collected every 5 min. Still images corresponding to this video are displayed in **Figure 2**. Scale bar is 20 µm.

**Video 3.** Time-lapse confocal microscopy of *P. palmivora* LILI-td-NT hyphae differentiating haustoria during *N. benthamiana* leaf infection. Plastids autofluorescence is shown in magenta. Frames were collected every 6.6 s. Still images corresponding to this video are shown in **Figure 3**. Scale bar is 20 µm.

**Video 4.** Time-lapse confocal microscopy of *P. palmivora* LILI-td-NT hyphae during haustoria differentiation in *N. benthamiana* leaf tissues. Plastids autofluorescence is shown in magenta. Frames were collected every 4 s. Still images corresponding to this video are shown in **Figure 3**. Scale bar is 10 µm.

**Video 5.** Time-lapse confocal microscopy of a *P. palmivora* LILI-td-NT hyphal tip on V8 agar plate. Frames were collected every 4.5 s. Still images corresponding to this video are displayed in **Figure 3**. Scale bar is 20 µm.

**Video 6.** Time-lapse confocal microscopy of a *P. palmivora* LILI-td-NT hyphal tip on V8 agar plate. Frames were collected every 4.8 s. Still images corresponding to this video are displayed in **Figure 3**. Scale bar is 20 µm.

**Video 7.** Time-lapse confocal microscopy of a *P. palmivora* LILI-td-NT hyphal branch point on V8 agar plate. Frames were collected every 15 s. Still images corresponding to this video are displayed in **Figure 5**. Scale bar is 20 µm.

**Video 8.** Time-lapse confocal microscopy of *P. palmivora* LILI-td-NT mycelium on V8 agar plate without (DMSO control) (Video 8A) or with 10 mg/L benomyl (Video 8B). Frames were collected every 12.8 s and 13 s, respectively. Still images corresponding to this video are displayed in **Figure 6**. Scale bar is 20 µm.

**Video 9.** Time-lapse confocal microscopy of *P. palmivora* mycelium from a transgenic strain expressing a mCitrine-Centrin2 reporter and a nuclear-localized mTFP1 (LILI-NT-Ce) on V8 agar plate. Frames were collected every 3 s. This video is linked to **Figure 7**. Scale bar is 20 µm.

## Supporting information

Supporting table 2 - Primers

## Acknowledgments

We are grateful to Dr. François Nédélec (SLCU, Cambridge) for helpful discussions. We thanks Dr. Lothar Kalmbach (SLCU, Cambridge) for providing mScarlet fluorescent reporter and Dr. Raymond Wightman (SLCU, Cambridge) for assistance with confocal microscopy. We are grateful to Dr. Philip Carella (SLCU, Cambridge) for proofreading the manuscript.

The authors declare no competing financial interests.

## Author contributions

E. Evangelisti conceived the experimental strategy, conducted experiments, acquired data, analyzed data and wrote the manuscript. L. Shenhav, T. Yunusov, M. Le Naour-Vernet and P. Rink conducted experiments, acquired data and analyzed data. S. Schornack acquired funding, conceived the experimental strategy, analyzed data and wrote the manuscript.

