## Supporting Figures and Table S1 for "Centrin-anchored hydrodynamic shape changes underpin active nuclear rerouting in branched hyphae of an oomycete pathogen"

#### Centrin-anchored active rerouting underpins nuclear distribution in branched hyphae of a plant pathogenic oomycete

#### Supporting figures

### Figure S1

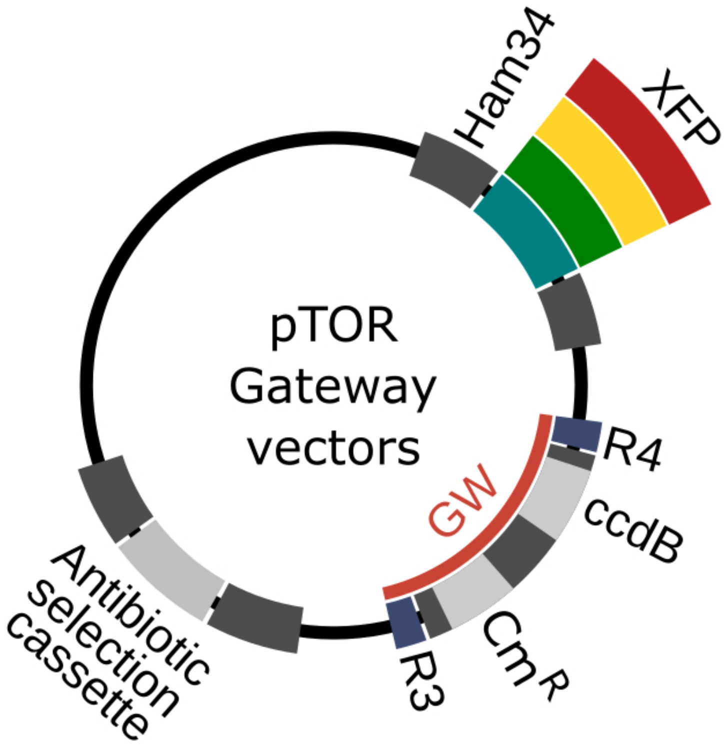

**Figure S1. Schematic representation of the pTOR-Gateway vector series.** A multisite attR4/attR3 Gateway insertion cassette allows for rapid testing of multiple promoter-reporter constructs. Adjacent is a cassette for constitutive expression of a cytoplasmic fluorescent reporter (either mTFP1, mWasabi, mCitrine or tdTomato) under the control of *Bremia lactucae* promoter Ham34. Vectors carry neomycin phosphotransferase (*nptII*) selectable marker.

### Figure S2

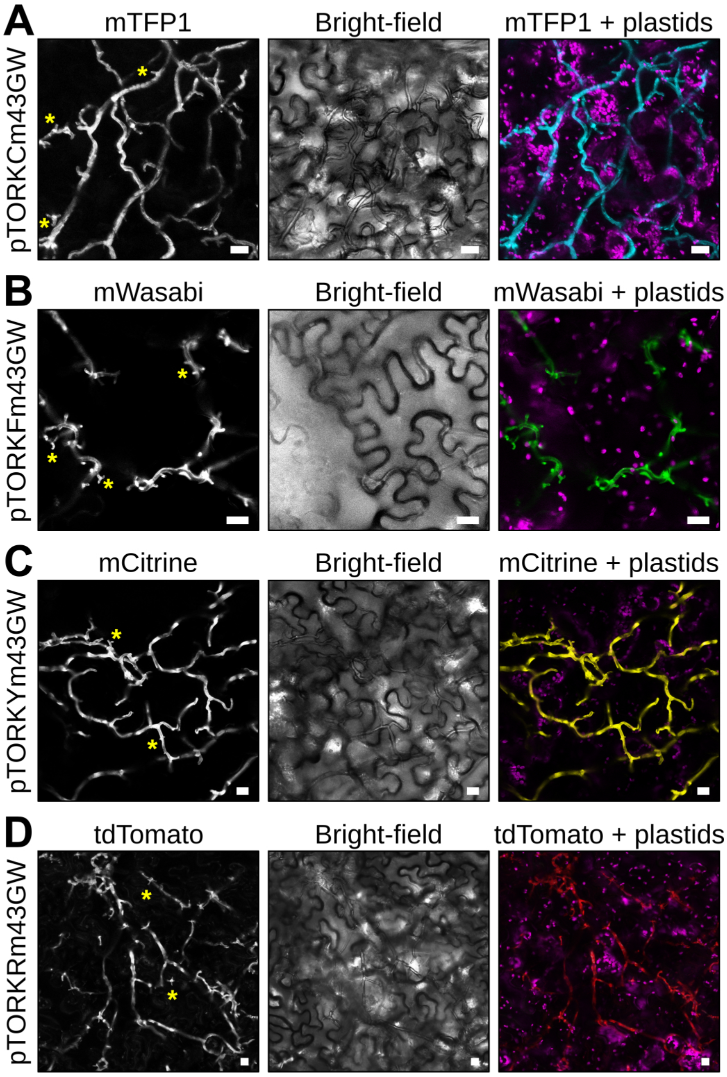

**Figure S2. Growth habit of transgenic *P. palmivora* strains carrying empty pTOR-Gateway vectors on *N. benthamiana* leaves.** (**A-D**) Leaves from 4-week-old *N. benthamiana* plants were inoculated with mycelium plugs of transgenic *P. palmivora* strains carrying empty pTOR-Gateway vectors. Fluorescence was monitored within leaf tissues after two days. Representative images of areas infected with mycelium expressing mTFP1 (**A**), mWasabi (**B**), mCitrine (**C**) and tdTomato (**D**) are shown. Yellow asterisks indicate haustoria. Scale bar is 10 μm.

### Figure S3

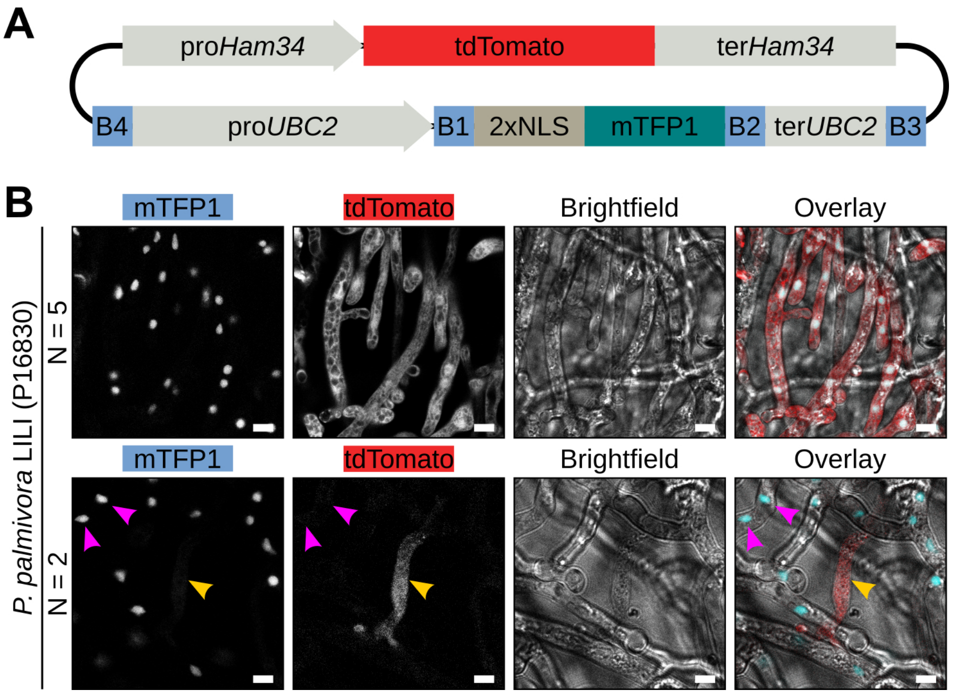

**Figure S3. Dual labelling of *P. palmivora* hyphae and nuclei.** **(A)** Schematic view of the construct used for dual labelling of nuclei and hyphae in *P. palmivora* strain LILI-td-NT. **(B)** Representative pictures of a transformant expressing tdTomato as well as a nuclear-localized mTFP1 (NLS:mTFP1) driven by the *P. palmivora ubiquitin-conjugating enzyme 2* (*UBC2*) native promoter. Representative pictures of transformants expressing both markers as well as rare cases of transformants expressing either tdTomato or mTFP1 in distinct hyphae. Scale bar is 10 µm.

### Figure S4

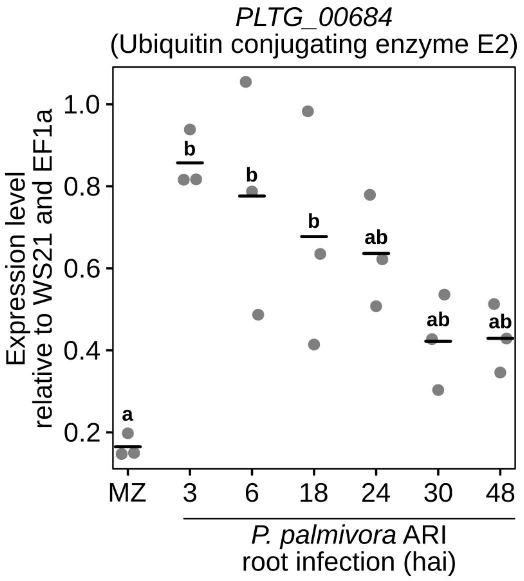

**Figure S4. *UBC2* transcript levels during *N. benthamiana* root infection.** *N. benthamiana* roots were inoculated with zoospores from the transgenic *P. palmivora* strain ARI-tdTomato (Le Fevre *et al*, 2016) and harvested at different times corresponding to early infection (3-6 hours), biotrophy (18-24 hai) and necrotrophy (30-48 hai). Expression data are given relative to *P. palmivora WS21* and *EF1α* reference genes. Statistical significance was assessed using one-way ANOVA and Tukey's HSD test ($P < 0.05$). MZ: axenically grown mycelium with sporangia.

### Figure S5

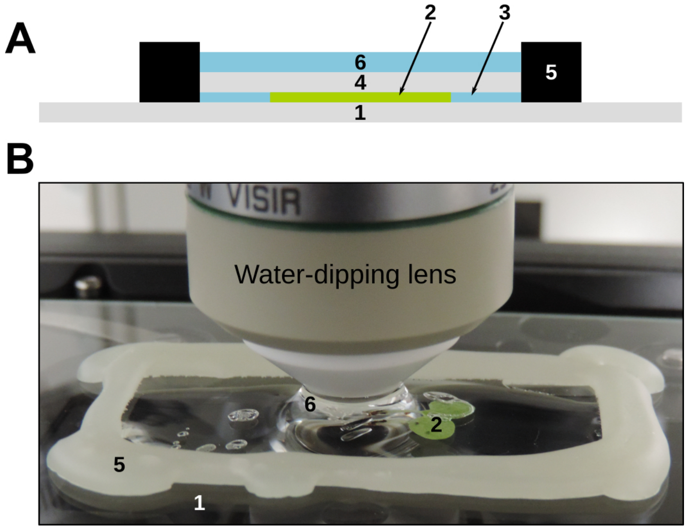

**Figure S5. Experimental set-up used for time-lapse imaging of infected *N. benthamiana* roots. (A)** Schematic representation of the experimental set-up. A one-week-old *N. benthamiana* seedling (2) is mounted in a liquid compartment (3) between a slide (1) and a coverslip (4). The edges of the coverslip are sealed with a 4:1 mix of paraffin and lanolin (5). The upper part of the coverslip is filled with water for use with a water-dipping objective. **(B)** Representative image of the experimental set-up. Numbers refer to the same elements as described previously.

### Figure S6

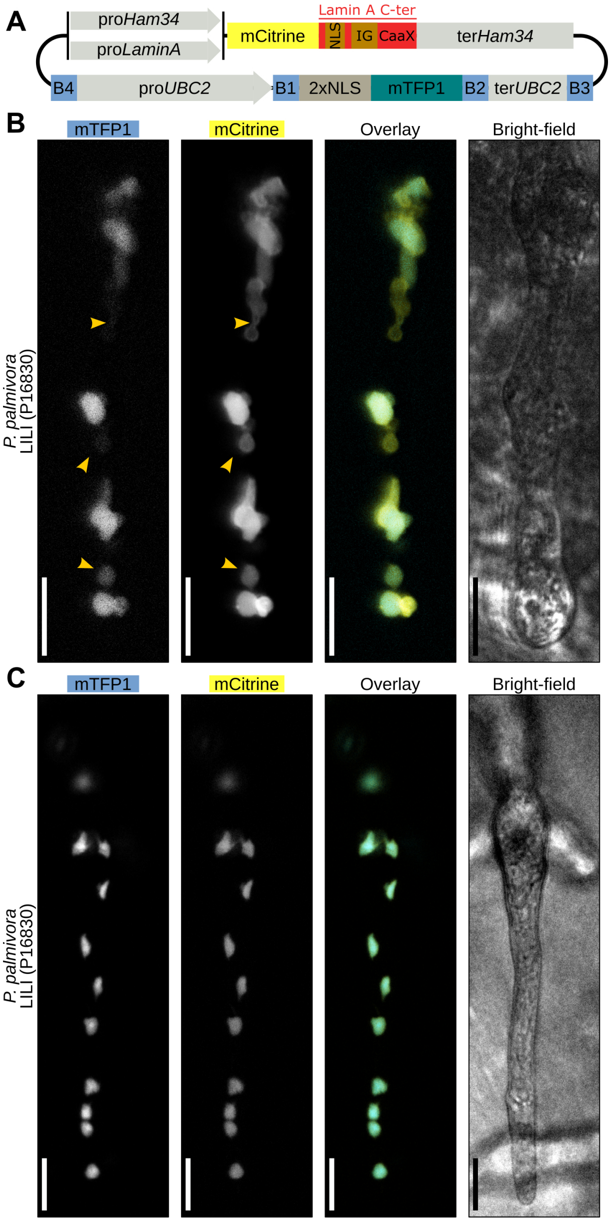

**Figure S6. Generation of a *P. palmivora* lamin A reporter.** (**A-C**) Transformation of *P. palmivora* LILI with a construct for constitutive (**A-B**) or native (**C-D**) expression of a mCitrine:LamA-Cter reporter. (**A**) Schematic view of the construct used for constitutive or native expression of the lamin reporter together with a nuclear-localized mTFP1. (**B**) Representative pictures of the bubbling phenotype observed upon constitutive expression of the lamin reporter. Arrowheads indicate bubbling nuclei. Scale bar is 10 µm. (**C**) Representative pictures of a hyphal segment upon native expression of the lamin reporter, showing absence of nuclear bubbling. Arrowheads indicate bubbling nuclei. Scale bar is 10 µm.

### Figure S7

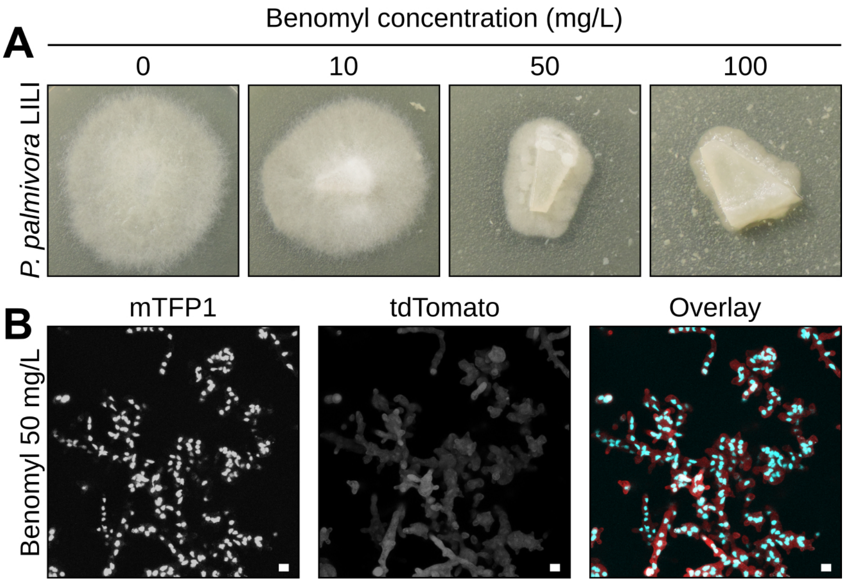

**Figure S7. Effect of antimicrotubule drug benomyl on *P. palmivora* growth.** (**A**) Representative pictures of *P. palmivora* LILI-td-NT mycelium growing on V8 agar plates supplemented or not with 10, 50 or 100 mg/L benomyl. (**B**) Confocal imaging of *P. palmivora* LILI-td-NT hyphae grown on V8 agar plates supplemented with 50 mg/L benomyl. Scale bar is 10 µm.

### Figure S8

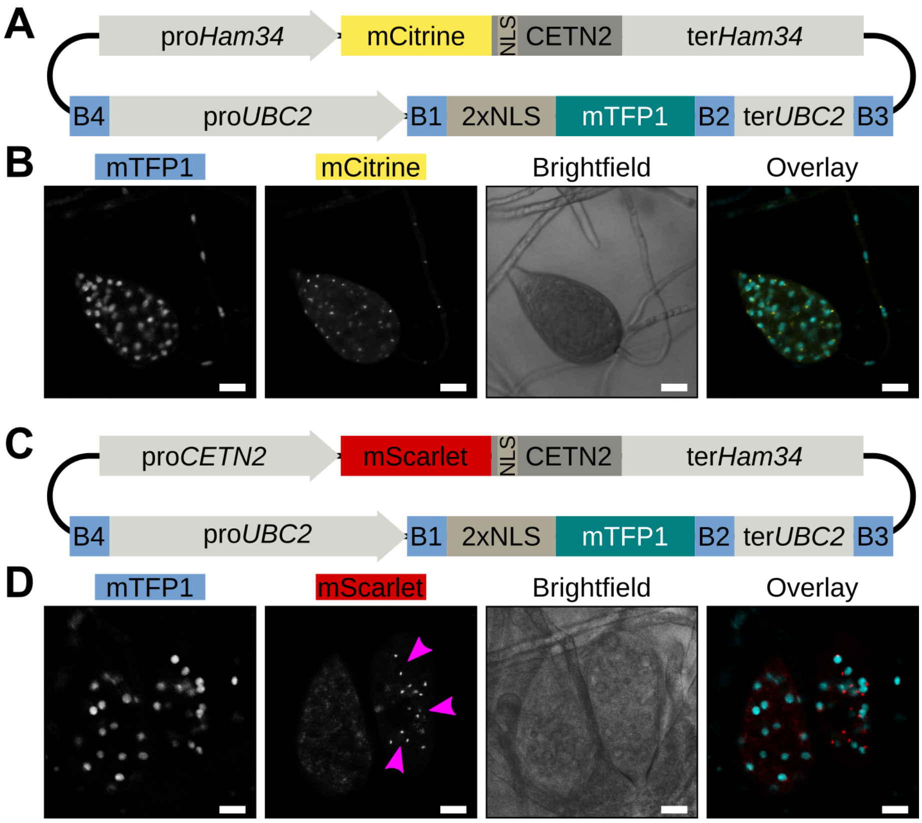

**Figure S8. Generation of a *P. palmivora* Centrin 2 (CETN2) reporter.** (**A**) Schematic view of the construct for Ham34-promoter-driven expression of a mCitrine:CETN2 reporter together with a nuclear-localized mTFP1. (**B**) Confocal imaging of a sporangium from a *P. palmivora* LILI-NT-Ce transgenics expressing the construct shown in (**A**). Scale bar is 10 µm. (**C**) Schematic view of the construct for CETN2-promoter-driven expression of a mScarlet:CETN2 reporter together with a constitutively expressed nuclear-localized mTFP1. (**D**) Confocal imaging of a sporangium from a transgenic *P. palmivora* strain expressing the construct shown in (**C**). Scale bar is 10 µm.

### Figure S9

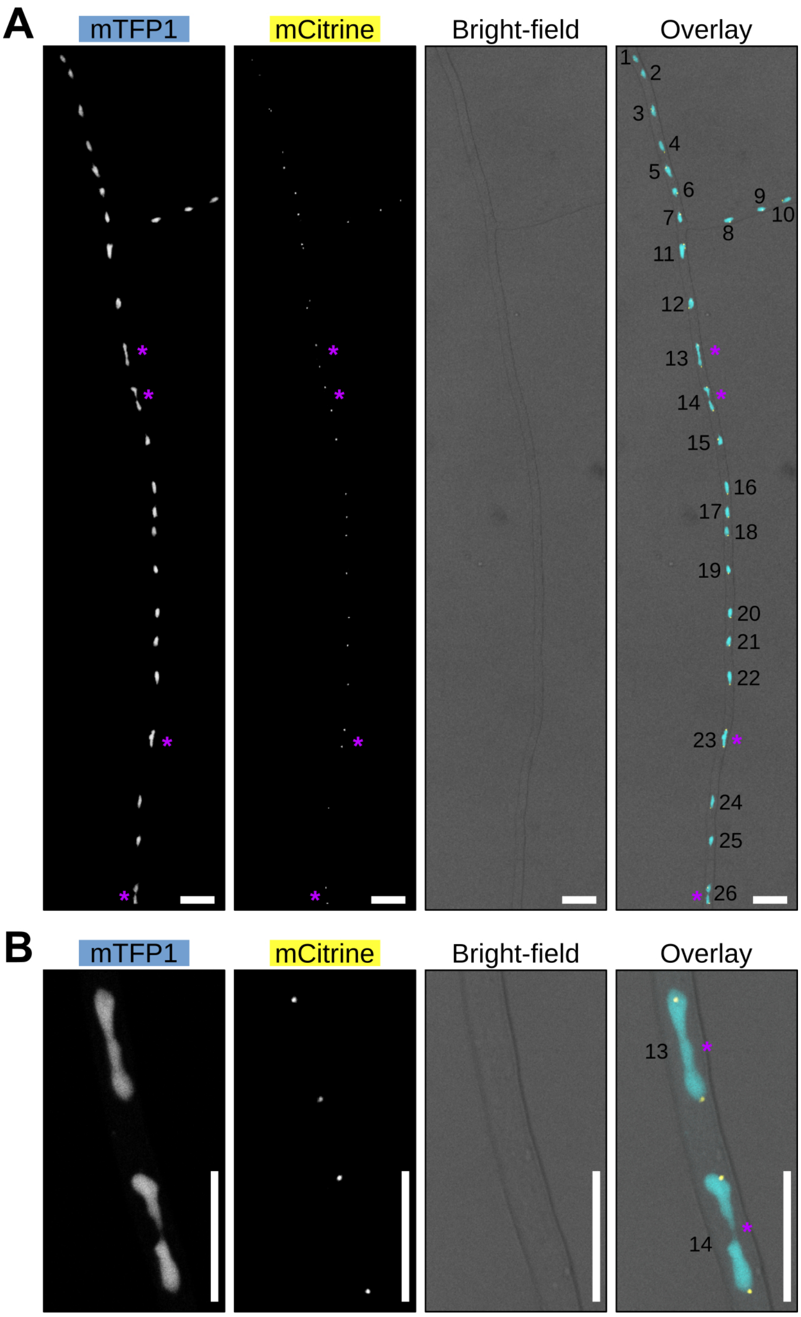

**Figure S9. Frequency of centrosome duplication within *P. palmivora* hyphae. (A-B)** Representative pictures of axenically-grown hyphae from the transgenic *P. palmivora* LILI-NT-Ce strain growing on V8 medium. **(A)** Distribution of nuclei and Centrin 2 (CETN2)-labelled centrosomes within an hyphal segment. Asterisks indicate nuclei with duplicated centrosomes. **(B)** Magnified views of nuclei 13 and 14. Scale bar is 10 µm.

#### Supporting tables

### Table S1

| Plasmid name | Fluorophore | Selection |
|---|---|---|
| pTOR**K**m43GW | *None* | G418 |
| pTOR**KC**m43GW | mTFP1 | G418 |
| pTOR**KF**m43GW | mWasabi | G418 |
| pTOR**KY**m43GW | mCitrine | G418 |
| pTOR**KR**m43GW | tdTomato | G418 |

**Table S1. pTOR-Gateway vectors**. pTOR-Gateway vectors follow naming conventions used for Gateway vectors. K indicates *nptII*, while C, F, Y and R stand for cyan, green, yellow and red fluorescence, respectively. The multi-site Gateway cassette carries attR4 and attR3 sites and hence was arbitrarily named m43GW in the absence of a T-DNA left border to define cassette orientation.

**Table S2**. Primers used in this study (xls file)

#### Supporting methods

**Quantitative reverse transcription-polymerase chain reaction (qRT-PCR) analyses**. Total RNA was extracted from axenically-grown *P. palmivora* mycelium containing sporangia (sample MZ) and infected *N. benthamiana* roots harvested at 3, 6, 18, 24, 30 and 48 h after inoculation (hai) using the RNeasy Plant Mini Kit (Qiagen, Germantown, USA). One microgram was reverse transcribed to generate first-strand complementary DNA (cDNA), using the Bio-Rad IScript cDNA Synthesis Kit according to the manufacturer's instructions (Bio-Rad, Hercules, USA). RNA quality was assessed by electrophoresis on agarose gel. Conditions for quantitative PCR were described previously (Evangelisti *et al*, 2017).
